# Hippocampal/medial temporal sclerosis is associated with clinical severity, regional TDP-43, and hippocampal atrophy in Alzheimer disease

**DOI:** 10.64898/2026.09.03.749099

**Authors:** Preeti Sharma, Rajath Devappa, Sivaprakasam R. Saroja

## Abstract

Alzheimer disease neuropathologic change (ADNC) does not fully explain clinical severity, suggesting important contributions from coexisting pathologies. We examined major co-pathologies in a multicenter autopsy cohort, focusing on hippocampal/medial temporal sclerosis (HS/MTL sclerosis), clinical severity, regional TDP-43 topography, clinical status near death, and gross hippocampal atrophy. We analyzed longitudinal clinical and neuropathology data from the National Alzheimer’s Coordinating Center autopsy cohort. Form-specific definitions harmonized AD pathology, major co-pathologies, HS/MTL sclerosis, and regional TDP-43 findings across neuropathology versions 8–10. HS/MTL sclerosis was present in 13.0% of assessed AD cases and was associated with greater CDR® Dementia Staging Instrument sum-of-boxes severity in non-AD (odds ratio [OR], 4.93; 95% confidence interval [CI], 3.66–6.63) and AD strata (OR, 2.56; 95% CI, 2.15–3.04). In version 10, HS/MTL sclerosis was associated with higher regional TDP-43 topography (OR, 6.54; 95% CI, 3.88–11.02) and greater gross hippocampal atrophy (OR, 6.43; 95% CI, 4.97–8.32). Among AD cases assessed within 2 years of death, HS-negative participants had greater odds of non-dementia status than HS-positive participants (OR, 5.81; 95% CI, 2.80–12.06). HS/MTL sclerosis marked greater clinical severity in both AD and non-AD strata and co-occurred with regional TDP-43 involvement and hippocampal atrophy. Its absence identified a subset with intermediate/high AD pathology but no dementia near death. Clinical severity therefore cannot be inferred from ADNC alone.

## BACKGROUND

Alzheimer disease (AD) is defined neuropathologically by the burden and distribution of amyloid-β plaques, tau neurofibrillary tangles, and neuritic plaques. Individuals with comparable AD neuropathologic change (ADNC) can differ markedly in cognitive and functional impairment [1,2]. Lewy body pathology, vascular brain injury, cerebral amyloid angiopathy (CAA), TDP-43 pathology, and hippocampal sclerosis commonly coexist with ADNC and contribute to dementia in late life [3,4].

Hippocampal/medial temporal sclerosis (HS/MTL sclerosis) is characterized by disproportionate neuronal loss and gliosis in hippocampal CA1 and/or the subiculum. It is associated with memory impairment and late-life dementia and can mimic AD clinically [8,10,15]. Its close relationship to TDP-43 was recognized before limbic-predominant age-related TDP-43 encephalopathy neuropathologic change (LATE-NC) was defined, and subsequent studies have shown that HS of aging frequently accompanies regional TDP-43 involvement and vascular pathology [8-13]. NACC records HS/MTL sclerosis across neuropathology form versions and regional TDP-43 involvement in NP v10, but these fields cannot establish a formal LATE-NC diagnosis or stage. Current consensus criteria distinguish LATE-NC from FTLD-TDP, ALS-TDP, and other TDP-43 proteinopathies; however, NACC does not contain all of the neuropathologic information needed to make these distinctions [20].

Previous clinicopathological studies linked both HS and TDP-43 to dementia and memory loss, including in people with ADNC [8,10,14]. Conversely, clinical preservation despite substantial AD pathology is more often observed when additional non-AD lesions, including hippocampal sclerosis, are absent [16,18,19]. However, the clinical association of HS/MTL sclerosis in the presence of ADNC remains unclear. It is also unknown whether the absence of HS identifies a less impaired phenotype near death and whether HS-related clinical severity is accompanied by regional TDP-43 involvement and gross hippocampal atrophy.

We used the NACC autopsy cohort, which links standardized longitudinal clinical assessments to neuropathology records from multiple U.S. centers [5,6]. We first determined the frequency and clinical associations of major co-pathologies using form-specific definitions and AD-alone reference groups based on documented absence of the lesion under study. We then focused on HS/MTL sclerosis and examined its relationship with CDR-SB severity across AD strata, regional TDP-43 topography and memory, non-dementia status near death, and gross hippocampal atrophy. We hypothesized that HS would be associated with greater clinical severity in both AD and non-AD strata and with gross hippocampal atrophy. We further hypothesized that the absence of HS would be associated with non-dementia status near death among participants with intermediate/high AD pathology and an eligible assessment.

## METHODS

### Study design, data source, and cohorts

We analyzed clinical and neuropathology records from the multicenter NACC autopsy cohort, which collects standardized data across U.S. Alzheimer’s Disease Research Centers (ADRCs) and links longitudinal clinical assessments to autopsy findings [5,6]. This analysis included 29 ADRCs and neuropathology form versions 8, 9, and 10. The source file included 49,614 participants. Of these, 5,638 had a supported NP form version (8, 9, or 10), and 3,587 met the harmonized criteria for intermediate/high AD pathology. The primary clinical AD cohort required at least one NACCUDSD code 2, 3, or 4 during follow-up. The death-aligned cohort required at least two valid CDR-SB visits, a valid month and year of death, and no CDR-SB assessment after death (Supplementary Tables S1 and S2).

### Pathology definitions and reference groups

Intermediate/high AD pathology was defined as NPADNC=2/3 in NP v10 or NPNIT=1/2 in NP v8/9. Non-AD was defined as NPADNC=0/1 in v10 or NPNIT=3/4 in v8/9. For the broad and clinical-severity analyses, HS/MTL sclerosis was classified as NPHIPSCL=1/2/3 versus 0 in NP v10 and NPSCL=1 versus 2 in NP v8/9. The primary NP v10 topography analysis required resolved laterality and compared NPHIPSCL=1/2 with 0; code 3 was included only in the sensitivity analysis. Lewy pathology, infarct/lacune, microinfarct, hemorrhage/microbleed, CAA, arteriolosclerosis, and regional TDP-43 involvement were defined from the corresponding NACC neuropathology fields. The Core AD-alone reference group required harmonized AD pathology together with documented assessment and absence of Lewy pathology, infarct/lacune, and HS/MTL sclerosis; CAA was allowed. The broad regional TDP-43 comparison used the strict NP v10 AD-alone reference group (Supplementary Table S1). Regional TDP-43 topography was classified as negative when NPTDPB-E were all negative; limbic-restricted when one or more NPTDPB-D fields were positive and NPTDPE was negative; or neocortical when NPTDPE was positive, regardless of the other fields. These groups describe regional distribution rather than burden, disease stage, or direction of spread. The available fields cannot distinguish LATE-NC from FTLD-TDP, ALS-TDP, or other TDP-43 disease contexts [21] (Supplementary Tables S1 and S3).

### Clinical, cognitive, and structural endpoints

CDR-SB scores were grouped as 0, 0.5-4, 4.5-9, 9.5-15.5, and 16-18. For death-aligned analyses, the date of death was assigned to the 15th day of the recorded month and year, and visits after that date were excluded. Logical Memory scores were harmonized only within supported form-version-by-language strata; raw scores from non-equivalent tests were not pooled. Gross hippocampal atrophy in NP v10 (NPGRHA) was categorized as none, mild, moderate, or severe after excluding codes 8, 9, -4, and blank values. Both NPGRHA and HS were recorded at autopsy.

For participants with harmonized intermediate/high AD pathology, we selected the valid visit closest to death that occurred on or before death and within 730.5 days. The primary endpoint was non-dementia clinical status near death, defined as NACCUDSD 1/2/3/8 versus 4. At the same visit, CDRGLOB 0/0.5 versus 1/2/3 provided a stricter sensitivity definition of lower clinical severity (Supplementary Table S4).

### Statistical analysis

We used clustered ordinal regression to examine CDR-SB severity. Each model included a cubic B-spline with four degrees of freedom for years before death, baseline age, sex, education, center, UDS/form version, remote assessment, and language. The focused HS model also included harmonized AD pathology and an AD×HS interaction. Inference used participant-clustered robust sandwich estimates. For participant-weighted marginal estimates, visits were first averaged within each participant and participants were then weighted equally. Each of the seven broad pathology models used a target-specific assessed cohort, comparator, and AD-alone reference group. The primary regional TDP-43 cohort required complete NPTDPB-E documentation and an HS assessment with resolved laterality. Adjusted topography models also required age at death, sex, education, and center. The primary model included these four covariates; the interaction model also included harmonized AD pathology and AD×HS. Logical Memory was modeled with flexible death-aligned time, HS, AD pathology, baseline age, sex, education, center, form version, and baseline clinical status. The limbic-restricted and neocortical contrasts formed a two-test Benjamini-Hochberg family. APOE ε4 dose was added in a paired sensitivity analysis restricted to participants with known genotype. Exploratory analyses examined hallucination symptoms and Trail Making Test Part B.

Near-death endpoints were analyzed using participant-level Firth logistic regression. Models included age at the selected visit, sex, education, center, UDS/form version, language, remote assessment, exact years before death, and the AD pathology category defined within each form version. The primary NP v10 ordinal gross-atrophy model included age at death, sex, education, center, categorical NPADNC, Lewy pathology, infarct/lacune, microinfarct, hemorrhage/microbleed, and CAA. Separate sensitivity analyses restricted HS to records with resolved laterality or added regional TDP-43 to examine pathological overlap. Paired complete-case sensitivity analyses added ordinal NACCARTE to the HS severity and gross-atrophy models. The joint death-aligned ordinal CDR-SB model included HS, NPGRHA, AD burden, and the same major measured co-pathologies; HS×atrophy and pathology×time interactions were not included. Benjamini-Hochberg correction was applied separately to the seven broad pathology main-effect tests, the two Logical Memory topography contrasts, and the two coefficients of interest in the joint HS-NPGRHA CDR-SB model.

## RESULTS

Table 1 summarizes the major analytic cohorts. Sample sizes differ across analyses because each denominator reflects the relevant eligibility criteria, availability of the pathology and outcome measures, and complete covariate data.

**Table 1.**
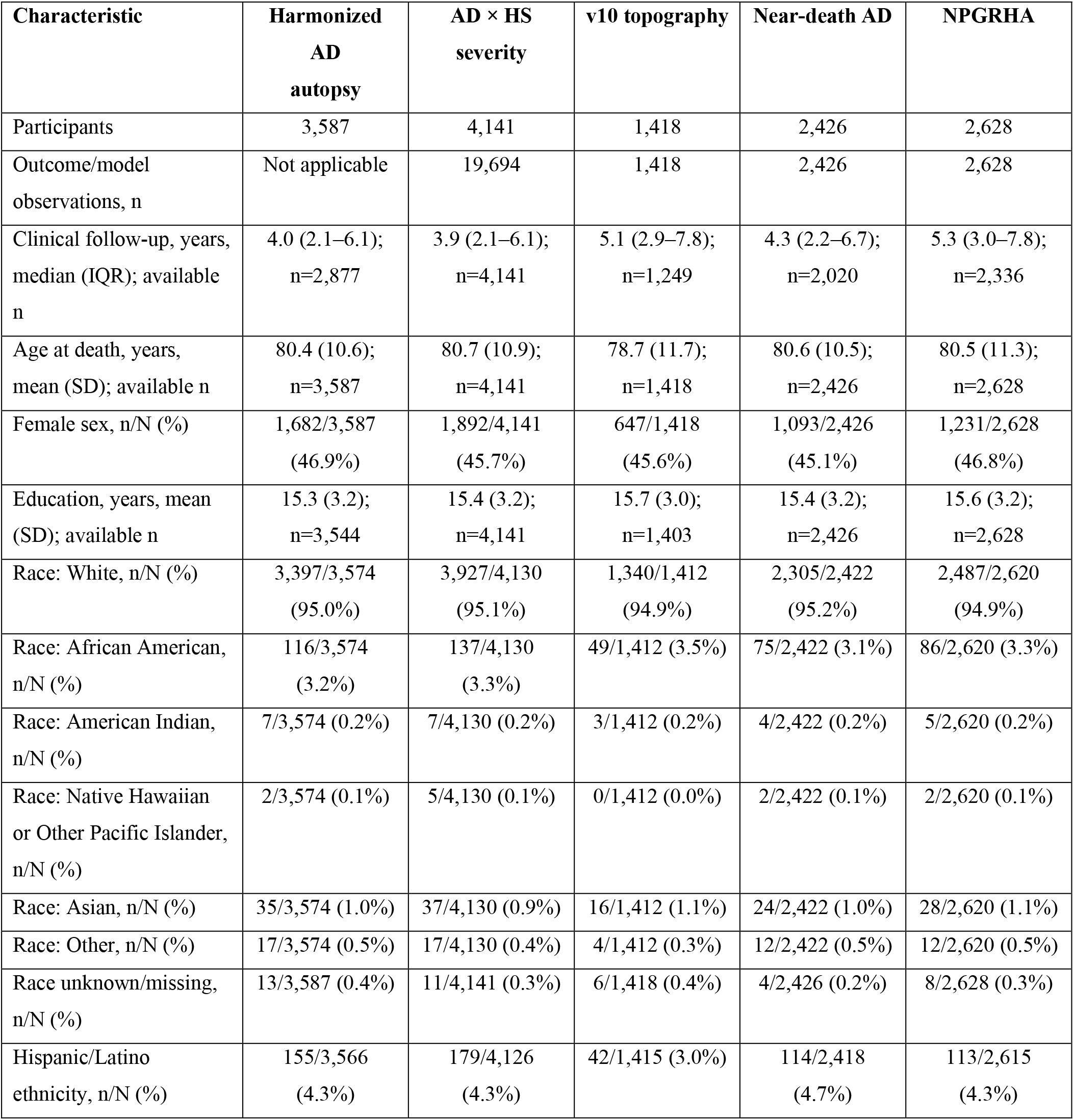
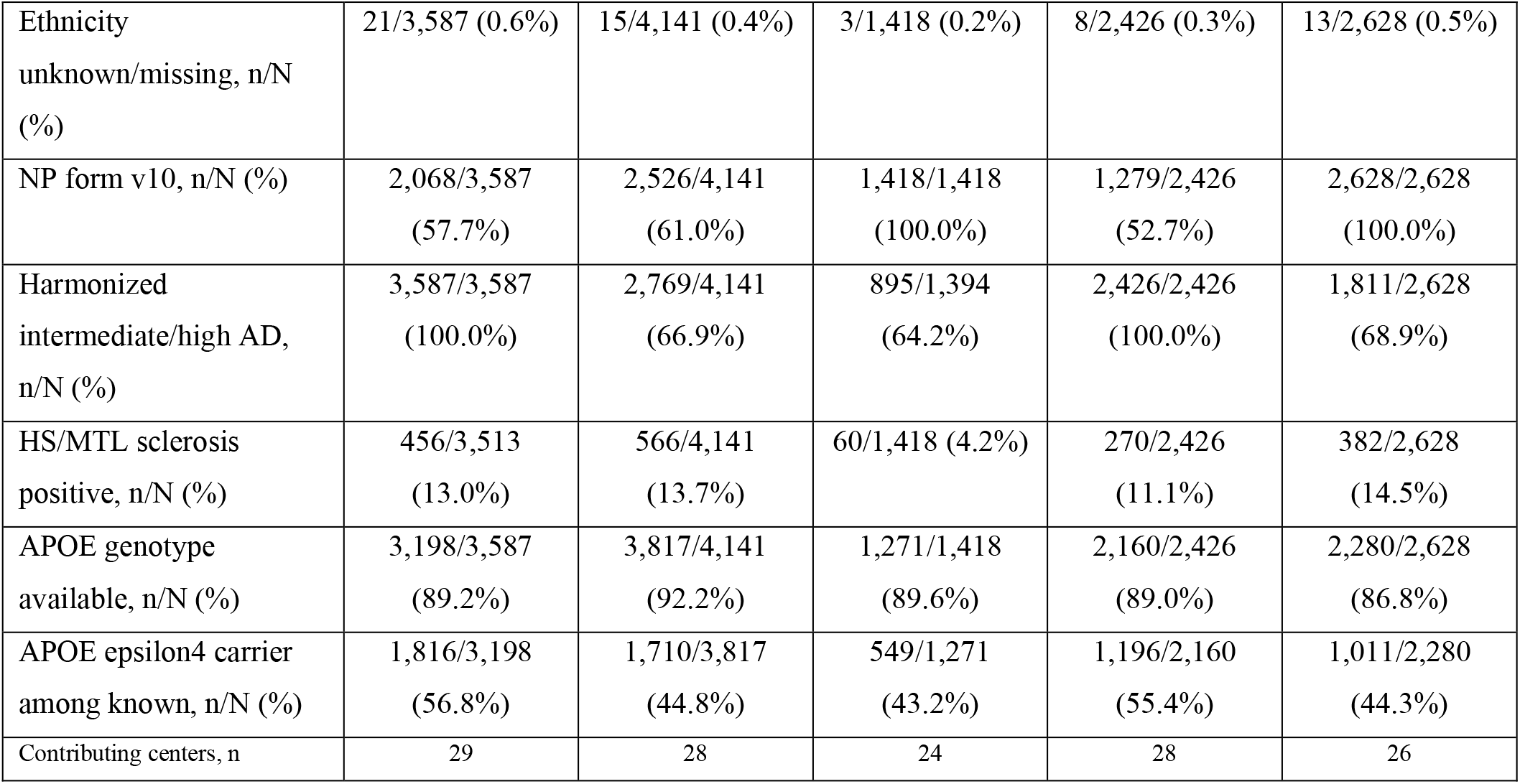
Descriptive characteristics of the major analytic cohorts.

### Co-pathologies in the autopsy-confirmed AD cohort

Before comparing co-pathologies, we established how the pathology and longitudinal-data requirements reduced the source cohort. The cohort included 49,614 participants, of whom 5,638 had supported NP v8/9/10 autopsy data. Among these participants, 3,587 met the harmonized criteria for intermediate/high AD pathology, 3,469 entered the primary clinical cohort, 2,776 had at least two valid CDR-SB observations, and 2,751 entered the death-aligned cohort (**Fig. 1A**). These nested cohorts provided the denominators for the broad and focused analyses that follow.

**Figure 1.**
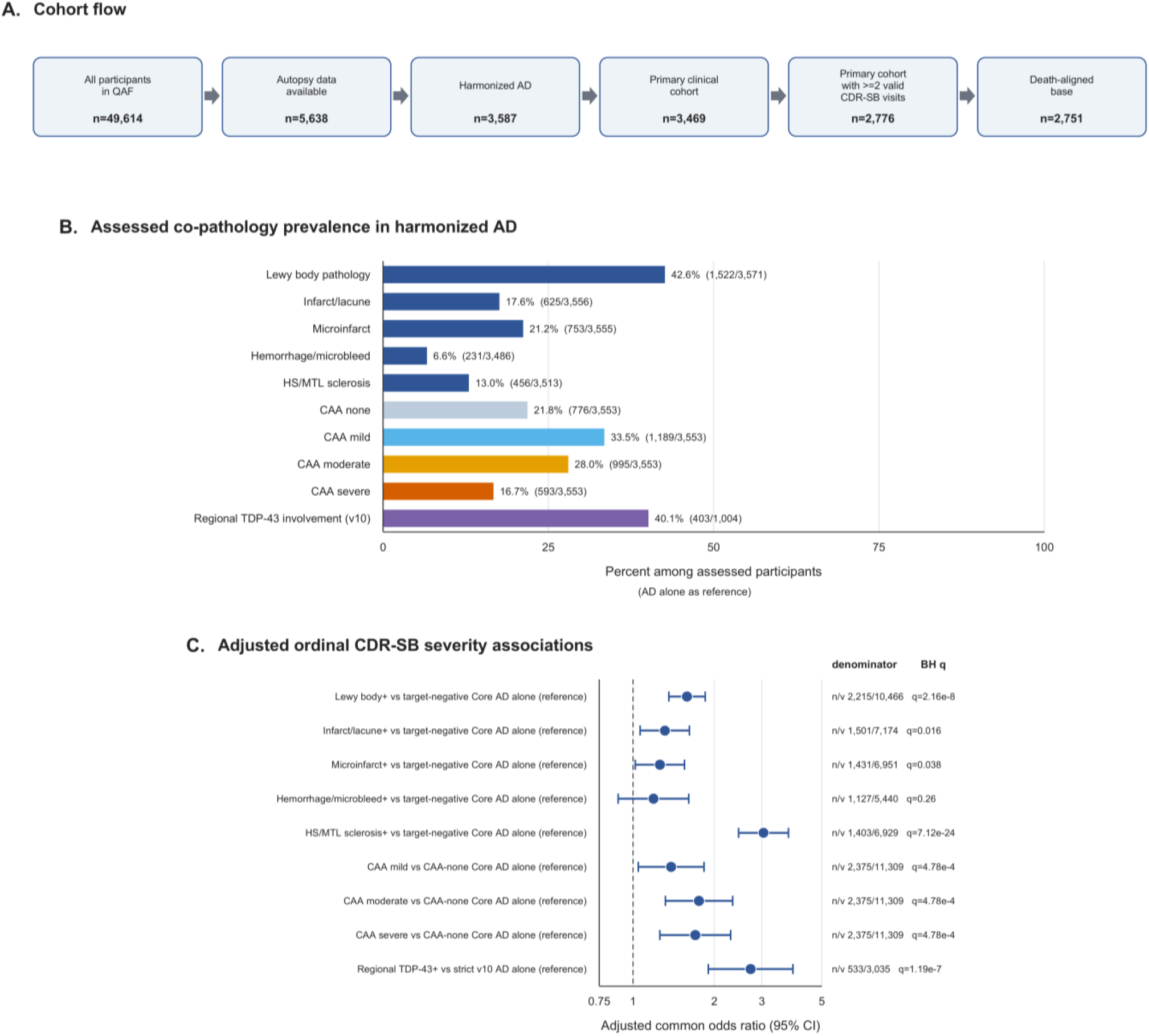
Co-pathologies in autopsy-confirmed AD. **A)** Cohort flow from the 49,614-participant QAF to 5,638 participants with supported NP v8/9/10 autopsy data, 3,587 with harmonized intermediate/high AD pathology, 3,469 in the primary clinical cohort, 2,776 with at least two valid CDR-SB visits, and 2,751 in the death-aligned cohort. **B)** Prevalence of each co-pathology among harmonized AD cases in whom that pathology was assessed. CAA is shown as the mutually exclusive categories none, mild, moderate, and severe. The Core AD-alone reference group included 1,408 participants, and the strict NP v10 AD-alone reference group included 204. **C)** Adjusted associations with ordinal CDR-SB severity. When proportional odds were not supported, common ORs are shown as overall summaries; threshold-specific estimates are provided in Supplementary Table S5. The n/v column reports participants/visits. The q values are BH-adjusted across the seven pathology tests, with a shared omnibus q value for the CAA categories.

Because lesion frequencies can only be interpreted among participants who were assessed, we calculated prevalence separately for each co-pathology. Co-pathologies were frequent among participants with harmonized AD pathology. Lewy body pathology was present in 1,522 of 3,571 participants (42.6%), infarct/lacune in 625 of 3,556 (17.6%), microinfarct in 753 of 3,555 (21.2%), hemorrhage/microbleed in 231 of 3,486 (6.6%), and HS/MTL sclerosis in 456 of 3,513 (13.0%). Mild, moderate, or severe CAA was recorded in 2,777 of 3,553 participants (78.2%). Regional TDP-43 involvement was present in 403 of 1,004 harmonized AD cases with the relevant v10 assessments (40.1%). The Core AD-alone reference group contained 1,408 participants, whereas the strict NP v10 AD-alone reference group contained 204 participants (**Fig. 1B**).

Version-specific AD definitions are provided in Supplementary Table S1. **Figure S1A** shows the assessed and unassessed denominators, and **Fig. S1B** shows the construction of the Core and strict NP v10 AD-alone reference groups. Thus, co-pathology was common in harmonized AD, whereas the strict v10 AD-alone reference group was relatively small. Prevalence alone does not show whether a lesion is clinically relevant. We therefore compared ordinal CDR-SB severity for each co-pathology with its target-specific AD-alone reference group. After correction across the seven pathology tests, Lewy pathology, infarct/lacune, microinfarct, HS/MTL sclerosis, CAA, and regional TDP-43 involvement were associated with greater ordinal CDR-SB severity. Hemorrhage/microbleed was not associated with severity in this analysis (**Fig. 1C**). Pairwise overlap among the measured co-pathologies was limited (**Fig. S2A**), and variance inflation factors did not indicate substantial collinearity (**Fig. S2B**). The broad screen therefore identified HS/MTL sclerosis and regional TDP-43 as clinically relevant features and provided the rationale for the focused analyses below.

### HS/MTL sclerosis and clinical severity in AD and non-AD strata

The broad screen showed a strong association between HS/MTL sclerosis and CDR-SB severity. We therefore examined whether this relationship differed according to the presence of intermediate/high AD pathology. The focused HS analysis included 4,141 participants and 19,694 visits. HS/MTL sclerosis was associated with greater ordinal CDR-SB severity in the non-AD stratum (OR, 4.93; 95% CI, 3.66-6.63; P=8.70×10^−26^) and the AD stratum (OR, 2.56; 95% CI, 2.15-3.04; P=1.23×10^−26^; **Fig. 2A**). The AD×HS interaction ratio was 0.52 (95% CI, 0.37-0.73; Pinteraction=0.000154), indicating a stronger relative association in non-AD than in AD. The association within AD nevertheless remained strong. Odds ratios do not show the absolute distribution of clinical severity. We therefore estimated participant-weighted CDR-SB category probabilities at fixed times before death. At -5, -3, and -1 years before death, participant-weighted estimates showed greater severity in participants with HS than in those without HS within each AD stratum (**Fig. 2B**). The adjusted distributions for non-AD participants with HS and AD participants without HS were also similar at each landmark. These distributions show that HS carries substantial clinical severity even outside the harmonized AD stratum.

**Figure 2.**
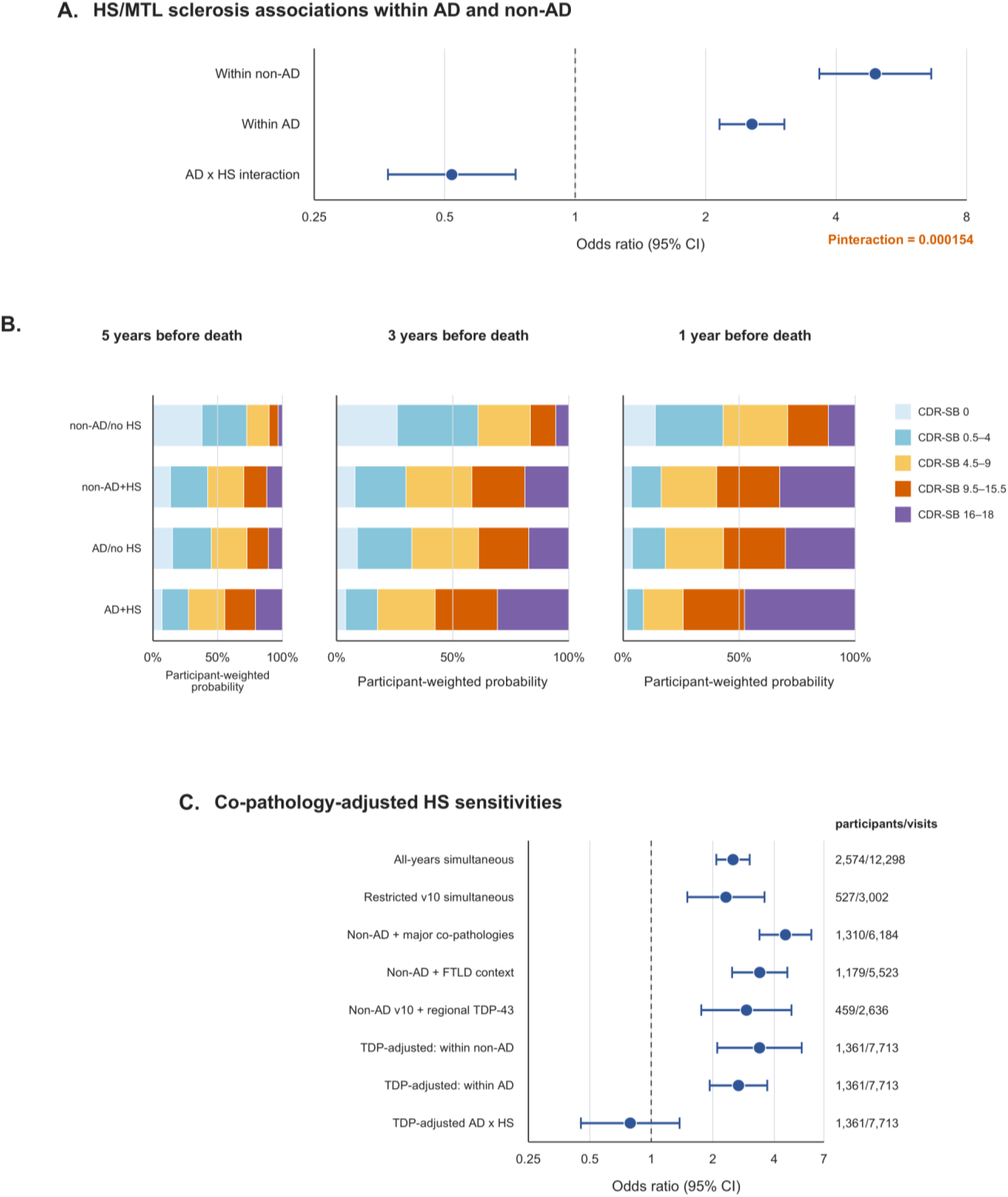
HS/MTL sclerosis and clinical severity. **A)** Adjusted associations between HS/MTL sclerosis and ordinal CDR-SB severity within autopsy-defined AD and non-AD strata, with the AD×HS interaction (Pinteraction = 0.000154). The AD stratum included all participants who met the harmonized AD definition, not only the AD-alone reference group. **B)** Participant-weighted adjusted CDR-SB distributions for non-AD/no HS, non-AD+HS, AD/no HS, and AD+HS at 5, 3, and 1 year before death. Visits were averaged within participants before participants were weighted equally. The model included no pathology×time interaction. **C)** Sensitivity analyses examined whether the HS associations changed after adjustment for major co-pathologies, restriction to NP v10, accounting for FTLD context, or adjustment for regional TDP-43 topography. The non-AD, AD, and AD×HS estimates were derived from the same TDP-complete cohort.

Because HS frequently occurs with other lesions, we next tested whether its association with severity persisted under increasingly stringent co-pathology adjustment. The association between HS/MTL sclerosis and severity remained after simultaneous adjustment for major measured co-pathologies (OR, 2.51; 95% CI, 2.08-3.03; BH q=4.64×10^−21^) and in the restricted v10 analysis (OR, 2.32; 95% CI, 1.50-3.58; BH q=0.00103; **Fig. 2C**). The corresponding conditional and fractional-logit analyses led to the same conclusion (**Fig. S2C-D**). Within the non-AD stratum, the corresponding ORs were 4.54 (95% CI, 3.39-6.08) after adjustment for major co-pathologies, 3.40 (95% CI, 2.49-4.64) after adjustment for broad FTLD context, and 2.92 (95% CI, 1.76-4.86) after adjustment for regional TDP-43 in the restricted v10 subset. With regional TDP-43 in the model, HS remained associated with severity in both strata, but the AD×HS interaction was attenuated (P=0.405). In the NACCARTE complete-case analysis, the ORs after adjustment for measured arteriolosclerosis were 4.61 in non-AD and 2.58 in AD, with an AD×HS ratio of 0.56 (**Fig. S2E-F**). HS therefore retained a strong association with severity across the sensitivity analyses.

### HS/MTL sclerosis, regional TDP-43 topography, and memory

HS/MTL sclerosis and regional TDP-43 were both associated with severity in the broad screen, raising the question of whether HS marks a distinct regional TDP-43 distribution. The primary NP v10 topography cohort included participants with complete HS and NPTDPB–E data and documented HS laterality. Among participants without HS/MTL sclerosis, 69% were TDP-43 negative, 23% had limbic-restricted involvement, and 8% had neocortical involvement. Among participants with HS/MTL sclerosis, 25% were TDP-43 negative, 42% had limbic-restricted involvement, and 33% had neocortical involvement (**Fig. 3A**). HS was therefore accompanied by a marked shift from TDP-43-negative status toward limbic and neocortical involvement.

**Figure 3.**
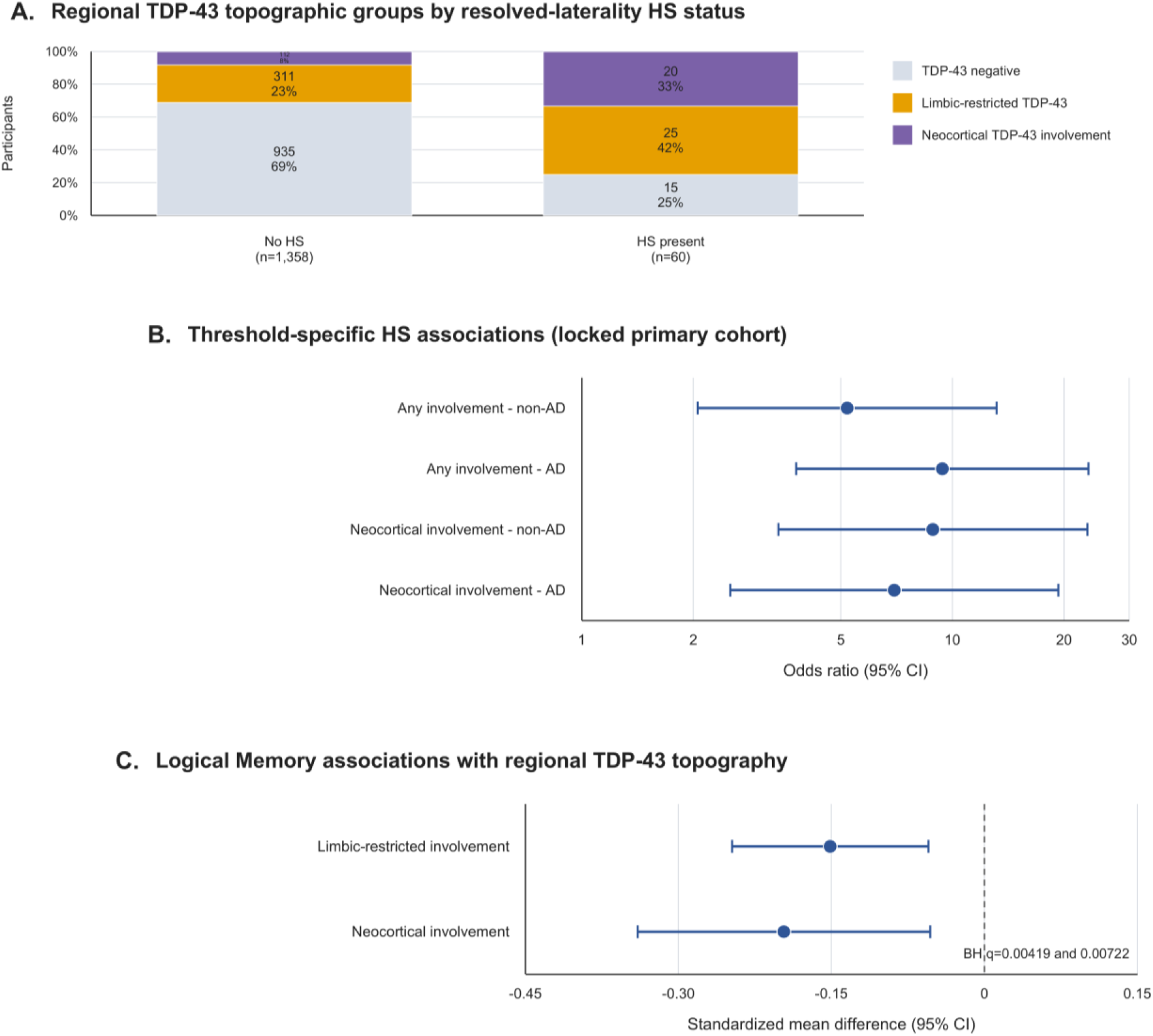
HS/MTL sclerosis and regional TDP-43 topography. **A)** Regional TDP-43 groups by HS status in the primary resolved-laterality NP v10 cohort (n=1,418; NPHIPSCL 1/2 versus 0; code 3 excluded). **B)** Threshold-specific HS associations in the covariate-complete primary cohort (n=1,379): any regional TDP-43 involvement versus negative, and neocortical involvement versus the lower topographic groups. The proportional-odds assumption was not met, so the common ordinal OR is not interpreted as uniform across thresholds. **C)** Adjusted Logical Memory associations for limbic-restricted and neocortical involvement relative to regional TDP-43 negativity (n=1,066; 3,701 visits), with BH correction across the two contrasts.

We next tested whether this shift persisted after covariate adjustment and whether it differed by AD status. HS/MTL sclerosis was associated with a higher regional TDP-43 topographic group in the common ordinal model (OR, 6.54; 95% CI, 3.88-11.02; P=1.66×10^−12^). The proportional-odds assumption was not met, so this OR is a global summary rather than an estimate assumed to be constant across thresholds. The proportional-odds diagnostic is shown in **Fig. S3A**. Threshold-specific models associated HS/MTL sclerosis with both any regional TDP-43 involvement and neocortical involvement in the non-AD and AD strata (**Fig. 3B**).

There was no evidence of AD×HS heterogeneity at either threshold. Including laterality-unresolved HS increased the descriptive cohort to 1,572 participants and the adjusted cohort to 1,529; the direction and magnitude of the associations were similar (**Fig. S3B**). Thus, HS tracked with broader regional TDP-43 involvement in both AD and non-AD strata.

To determine whether regional TDP-43 topography had a domain-specific clinical correlate, we modeled Logical Memory performance. The Logical Memory analysis accounted for form version and language and included 1,066 participants and 3,701 visits. Compared with participants without regional TDP-43 involvement, those with limbic-restricted involvement had lower Logical Memory performance (standardized mean difference, -0.151; 95% CI, -0.247 to -0.055; BH q=0.00419).

Neocortical involvement was also associated with lower performance (-0.197; 95% CI, -0.340 to - 0.053; BH q=0.00722; **Fig. 3C**). Adding APOE ε4 dose in the paired subset with known genotype did not materially change either estimate. Both regional patterns were therefore associated with worse episodic memory Analyses of hallucination symptoms showed no association (**Fig. S3D**). We did not interpret the positive neocortical Trail-B estimate because test completion and missingness differed substantially by clinical severity and TDP-43 topography (**Fig. S3C**).

### HS status and non-dementia clinical status near death

Some participants die with substantial AD pathology without having reached a dementia-level clinical status. We examined whether HS distinguished this clinicopathologic discordance. The near-death analysis was restricted to participants with harmonized intermediate/high AD pathology. Of 3,587 AD cases, 3,513 had an HS/MTL sclerosis assessment and 2,451 had an eligible visit. The primary complete-case model included 2,426 participants, and 2,353 remained after adding the major co-pathology covariates (**Fig. 4A**). Non-dementia clinical status near death was defined before outcome modeling; a stricter CDRGLOB definition provided the sensitivity endpoint. The timing of the selected visits relative to death is shown in **Fig. S4A**. This approach placed the clinical assessment close to death while preserving the prespecified alternative endpoint.

**Figure 4.**
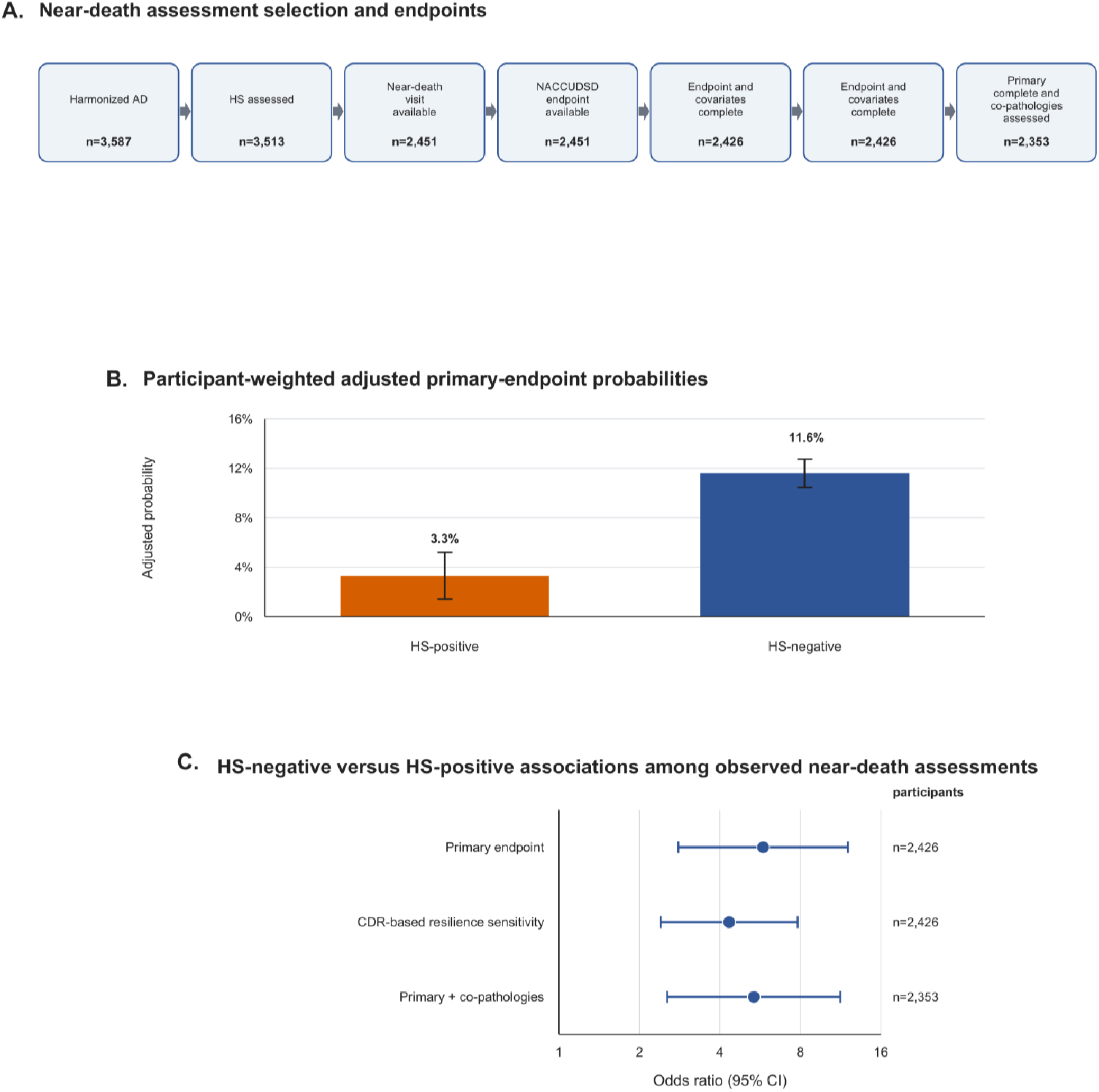
HS and non-dementia clinical status near death in AD. **A)** Cohort selection and definition of the two endpoints among participants with harmonized intermediate/high AD pathology. The primary endpoint was non-dementia clinical status, defined as NACCUDSD 1/2/3/8 versus 4. The stricter CDRGLOB sensitivity used 0/0.5 versus 1/2/3 at the same selected visit. B) Participant-weighted adjusted probabilities of the primary endpoint by HS status (n = 2,426). **C)** Associations comparing HS-negative with HS-positive participants for the primary endpoint, the CDRGLOB sensitivity, and the model adjusted for major co-pathologies.

We first estimated the absolute probability of non-dementia clinical status near death by HS status. The adjusted probabilities were 11.6% and 3.3%, respectively (**Fig. 4B**). Thus, non-dementia status near death was uncommon in both groups but was substantially less frequent when HS was present.

We then quantified this difference and tested its robustness to the stricter endpoint and additional co-pathology adjustment. HS-negative participants had greater odds of non-dementia clinical status near death than HS-positive participants (OR, 5.81; 95% CI, 2.80-12.06; P=2.36×10^−6^; **Fig. 4C**). The same pattern was observed with the stricter CDRGLOB definition (OR, 4.33; 95% CI, 2.40-7.81; P=1.13×10^−6^) and after adjustment for major measured co-pathologies (OR, 5.36; 95% CI, 2.54-11.30; P=1.01×10^−5^). Sparse nuisance levels and model calibration are summarized in **Fig. S4B-C**.

Across definitions and adjustments, the absence of HS was consistently associated with greater odds of preserved non-dementia status near death among participants with an observed near-death assessment.

### HS/MTL sclerosis, gross hippocampal atrophy, and CDR-SB severity

HS is defined microscopically. We therefore asked whether it corresponded to a gross structural change in the hippocampus. The primary gross-atrophy analysis included 2,628 participants with NP v10 data, of whom 382 were HS-positive. The adjusted probability of severe gross hippocampal atrophy was 47.6% in HS-positive participants and 16.0% in HS-negative participants (**Fig. 5A**). HS was therefore accompanied by a pronounced shift toward more severe gross hippocampal atrophy.

**Figure 5.**
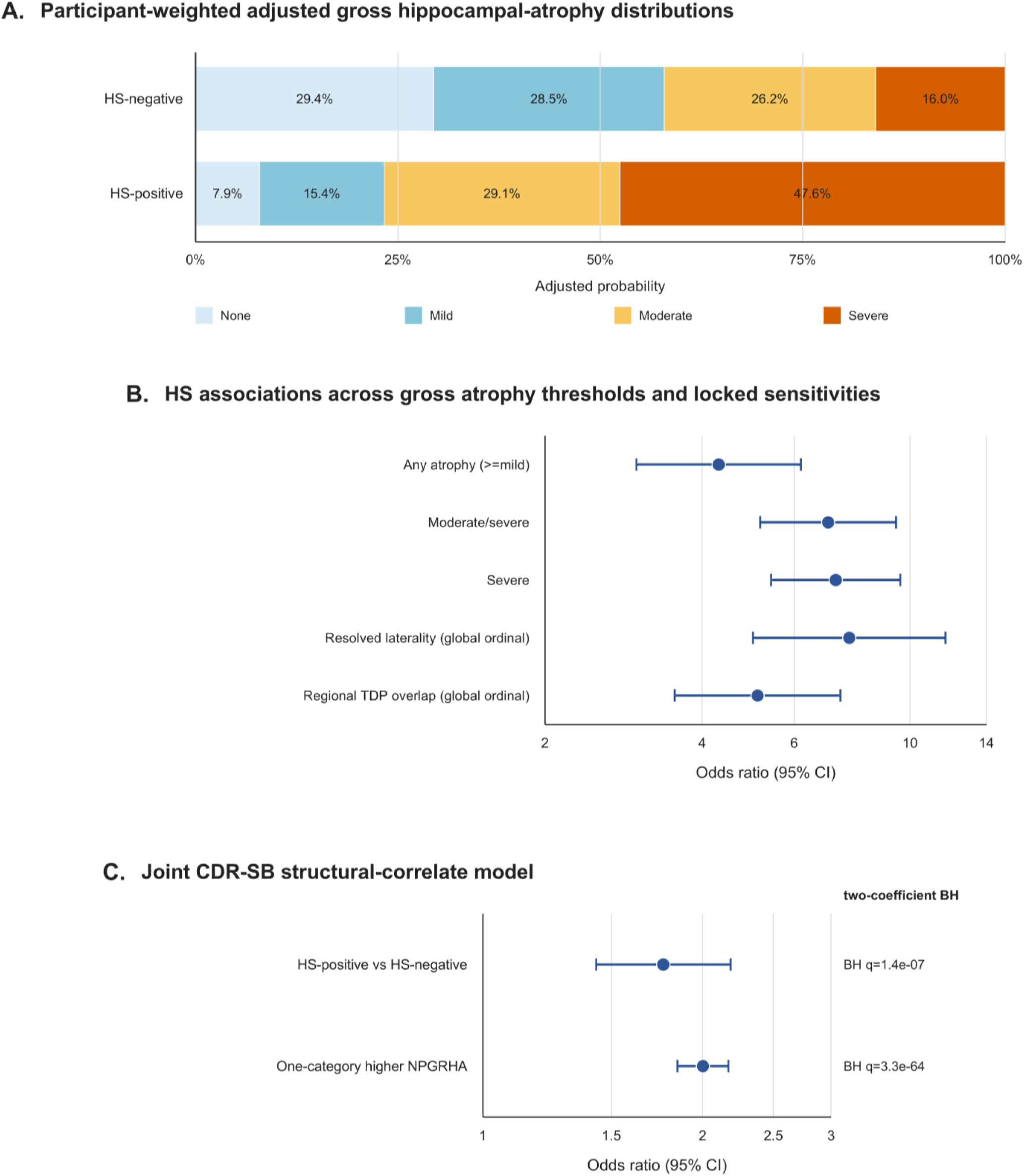
HS/MTL sclerosis and gross hippocampal atrophy. **A)** Participant-weighted adjusted NPGRHA distributions by HS status in the primary NP v10 complete-case cohort (n = 2,628). **B)** Threshold-specific associations between HS and any, moderate/severe, and severe gross hippocampal atrophy, with global ordinal sensitivity estimates after restricting HS to resolved laterality or adjusting for regional TDP-43 topography. Proportional odds were not supported (P = 0.00447); interpretation therefore rests on the threshold-specific estimates. **C)** Mutually adjusted associations of HS and a one-category increase in NPGRHA with ordinal CDR-SB severity (n = 2,315; 13,185 visits), with BH correction across the two coefficients. 00000

Because the proportional-odds assumption was not met, we examined the association across individual atrophy thresholds as well as with a global ordinal estimate. HS/MTL sclerosis was associated with a higher gross hippocampal atrophy category in the adjusted ordinal model (common OR, 6.43; 95% CI, 4.97-8.32; P=1.64×10^−45^; **Fig. 5B**). Because the proportional-odds assumption was not met (P=0.00447), the common OR is a global summary. The diagnostic is shown in **Fig. S5A**. Threshold-specific ORs were 4.30 (95% CI, 2.99-6.18) for any atrophy, 6.97 (95% CI, 5.16-9.40) for moderate/severe atrophy, and 7.20 (95% CI, 5.42-9.57) for severe atrophy. Restriction to records with resolved HS laterality gave a global OR of 7.65 (95% CI, 5.01-11.69); adding regional TDP-43 topography in the complete v10 subset gave a global OR of 5.11 (95% CI, 3.54-7.36). These sensitivity estimates are shown in **Fig. 5B**. In the NACCARTE-complete analysis, adjustment for measured arteriolosclerosis gave a common OR of 6.30; threshold-specific estimates and adjusted distributions are shown in **Fig. S5B-C**. The association between HS and gross hippocampal atrophy was therefore present across thresholds and remained after the laterality, TDP-43, and arteriolosclerosis sensitivity analyses.

Finally, we tested whether HS and gross hippocampal atrophy each retained an association with CDR-SB when modeled together. The joint death-aligned CDR-SB model included 2,315 participants and 13,185 visits. Both HS/MTL sclerosis (OR, 1.77; BH q=1.39×10^−7^) and each one-category increase in NPGRHA (OR, 2.00; BH q=3.33×10^−6^4) were associated with greater ordinal CDR-SB severity (**Fig. 5C**). These results indicate that HS and gross atrophy carry partly nonredundant associations with clinical severity; because both measures were obtained at autopsy, the model does not establish mediation or temporal direction.

## DISCUSSION

In the NACC autopsy cohort, HS/MTL sclerosis was strongly associated with greater CDR-SB severity in participants with and without AD pathology. The association remained after adjustment for major measured co-pathologies and arteriolosclerosis. HS/MTL sclerosis also co-occurred with limbic-restricted or neocortical TDP-43 involvement and marked hippocampal atrophy; the two TDP-43-positive groups had lower memory scores. In contrast, participants without HS/MTL sclerosis were more likely to have non-dementia clinical status near death despite intermediate/high AD pathology. ADNC alone therefore did not account for the clinical variation observed during life.

Co-pathology is common in AD. Lewy body pathology, vascular lesions, CAA, HS/MTL sclerosis, and regional TDP-43 involvement were each associated with greater CDR-SB severity in their respective analyses, consistent with earlier community and autopsy studies [3,4,7]. These estimates cannot be used to rank the lesions because the assessed denominator, reference group, and model differed for each analysis. We therefore focused HS analyses instead show that one lesion can be examined across AD strata, regional TDP-43 findings, gross hippocampal atrophy, and near-death clinical status.

HS/MTL sclerosis was strongly associated with regional TDP-43 involvement. TDP-43 pathology has long been observed in hippocampal sclerosis and AD [8,10], and LATE-NC is now recognized as an important contributor to late-life cognitive impairment [9,12,14]. Recent NACC analyses have also linked LATE-NC stage with HS of aging, hippocampal atrophy, and cognition [12]. In the present cohort, regional TDP-43 involvement was more common in participants with HS/MTL sclerosis, and both limbic-restricted and neocortical involvement were associated with lower Logical Memory scores. The regional NACC fields cannot measure TDP-43 burden, assign a formal LATE-NC stage, determine the direction of spread, or distinguish LATE-NC from FTLD-TDP, ALS-TDP, and other TDP-43 disease contexts. These cross-sectional autopsy data also cannot determine whether HS and TDP-43 arise from the same process.

Among participants with intermediate/high AD pathology and an eligible assessment near death, those without HS/MTL sclerosis were more likely to have non-dementia clinical status. The association remained under the stricter CDRGLOB definition and after adjustment for major measured co-pathologies. This finding identifies a less clinically impaired subset of participants with substantial AD pathology, consistent with earlier reports that clinical preservation is more frequent when additional lesions are absent [16,18,19]. It does not show that absence of HS protects against dementia. Because eligible near-death assessments were less common in HS-positive participants, the result is conditional on assessment availability and cannot estimate the frequency of this phenotype in all people with AD pathology.

HS/MTL sclerosis was also strongly associated with gross hippocampal atrophy. The associations with moderate/severe and severe atrophy remained after adjustment for AD burden, Lewy pathology, vascular lesions, and CAA. In the joint model, HS/MTL sclerosis and gross atrophy category each retained an association with greater CDR-SB severity. Sclerosis and atrophy were both recorded at autopsy, so their temporal order cannot be determined. Longitudinal MRI combined with fluid biomarkers will be needed to establish when each abnormality develops and whether they arise from a common TDP-43-associated process.

The multicenter sample, form-specific pathology definitions, lesion-specific assessed denominators, and documented-absence reference groups reduced several common sources of misclassification. The analysis plan defined the multiplicity families in advance and included sensitivity analyses for regional TDP-43, APOE availability, HS laterality coding, and measured arteriolosclerosis. We kept the broad co-pathology comparisons separate from the focused HS analyses because the lesions differed in assessment, prevalence, and biological meaning.

### Limitations and future directions

Neuropathology was measured only at death, whereas the clinical observations came from earlier visits. The data therefore cannot establish when HS, TDP-43 involvement, or atrophy developed, or whether one lesion mediated another. Independent autopsy cohorts with formal LATE-NC adjudication are needed to test the reproducibility and specificity of the associations. Longitudinal structural MRI linked to CSF or plasma markers of amyloid, tau, and neurodegeneration could determine when hippocampal atrophy emerges relative to clinical decline. Regional molecular and single-nucleus studies comparing AD alone, AD with HS/MTL sclerosis, and AD with regional TDP-43 involvement could then test whether the groups share a biological mechanism.

Across AD and non-AD strata, HS/MTL sclerosis marked greater clinical severity, regional TDP-43 involvement, and gross hippocampal atrophy. Its absence was associated with non-dementia status near death despite intermediate/high AD pathology. These results show the contribution of co-pathology to clinical heterogeneity but do not establish causation.

## Acknowledgments

The authors thank the National Alzheimer’s Coordinating Center and the participating Alzheimer’s Disease Research Centers for making the clinical and neuropathology data available. The NACC database is funded by NIA/NIH Grant U24 AG072122. NACC data are contributed by the NIA-funded ADRCs: P30 AG062429 (PI James Brewer, MD, PhD), P30 AG066468 (PI Oscar Lopez, MD), P30 AG062421 (PI Bradley Hyman, MD, PhD), P30 AG066509 (PI Thomas Grabowski, MD), P30 AG066514 (PI Mary Sano, PhD), P30 AG066530 (PI Helena Chui, MD), P30 AG066507 (PI Marilyn Albert, PhD), P30 AG066444 (PI David Holtzman, MD), P30 AG066518 (PI Lisa Silbert, MD, MCR), P30 AG066512 (PI Thomas Wisniewski, MD), P30 AG066462 (PI Scott Small, MD), P30 AG072979 (PI David Wolk, MD), P30 AG072972 (PI Charles DeCarli, MD), P30 AG072976 (PI Andrew Saykin, PsyD), P30 AG072975 (PI Julie A. Schneider, MD, MS), P30 AG072978 (PI Ann McKee, MD), P30 AG072977 (PI Robert Vassar, PhD), P30 AG066519 (PI Frank LaFerla, PhD), P30 AG062677 (PI Ronald Petersen, MD, PhD), P30 AG079280 (PI Jessica Langbaum, PhD), P30 AG062422 (PI Gil Rabinovici, MD), P30 AG066511 (PI Allan Levey, MD, PhD), P30 AG072946 (PI Linda Van Eldik, PhD), P30 AG062715 (PI Sanjay Asthana, MD, FRCP), P30 AG072973 (PI Russell Swerdlow, MD), P30 AG066506 (PI Glenn Smith, PhD, ABPP), P30 AG066508 (PI Stephen Strittmatter, MD, PhD), P30 AG066515 (PI Victor Henderson, MD, MS), P30 AG072947 (PI Suzanne Craft, PhD), P30 AG079231 (PI Henry Paulson, MD, PhD), P30 AG066546 (PI Sudha Seshadri, MD), P30 AG086401 (PI Erik Roberson, MD, PhD), P30 AG086404 (PI Gary Rosenberg, MD), P20 AG068082 (PI Angela Jefferson, PhD), P30 AG072958 (PI Heather Whitson, MD), P30 AG072959 (PI James Leverenz, MD).

## Author contributions

SRS, PS and RD contributed to data curation, analysis, interpretation, and revision of the manuscript. SRS conceived and supervised the study and drafted the manuscript. All authors reviewed and approved the final version.

## Ethics, consent, and data-use statement

This study used deidentified NACC data obtained under a data-use agreement. The contributing ADRCs obtained local institutional review board approval and written informed consent from participants or their legally authorized representatives. The present secondary analysis involved no direct participant contact.

## Data availability

NACC data are available to qualified investigators through the NACC data-request process under a data-use agreement. Participant-level data cannot be redistributed with this article.

---

**Supplementary figure S1.**
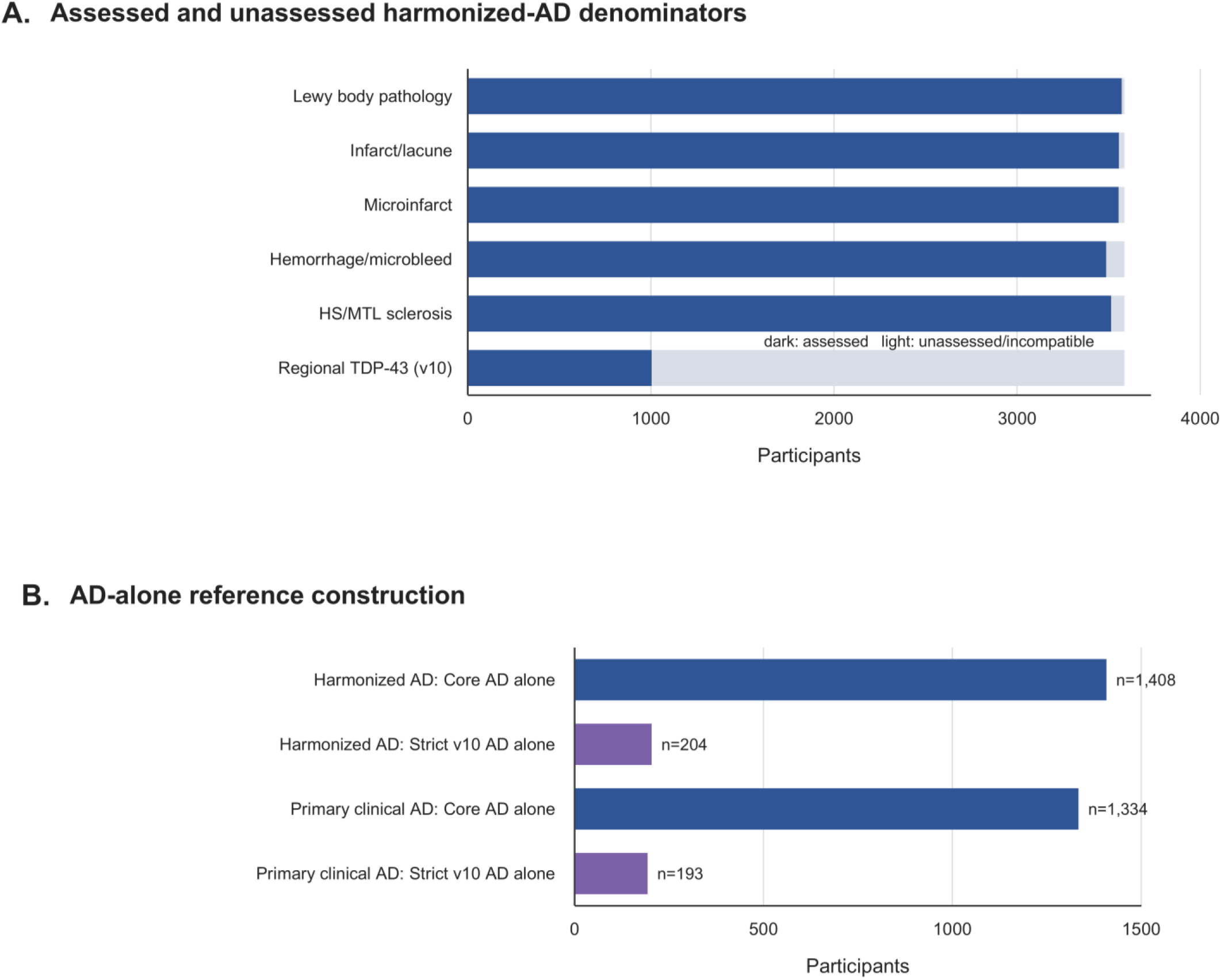
Pathology harmonization and construction of AD-alone reference groups. **A)** Assessed and unassessed denominators for each co-pathology among harmonized AD participants. **B)** Construction of the Core and strict NP v10 AD-alone reference groups in the harmonized AD and primary clinical cohorts. Unassessed, indeterminate, blank, or version-incompatible records were excluded from the applicable denominator and were not classified as pathology absent.

**Supplementary figure S2.**
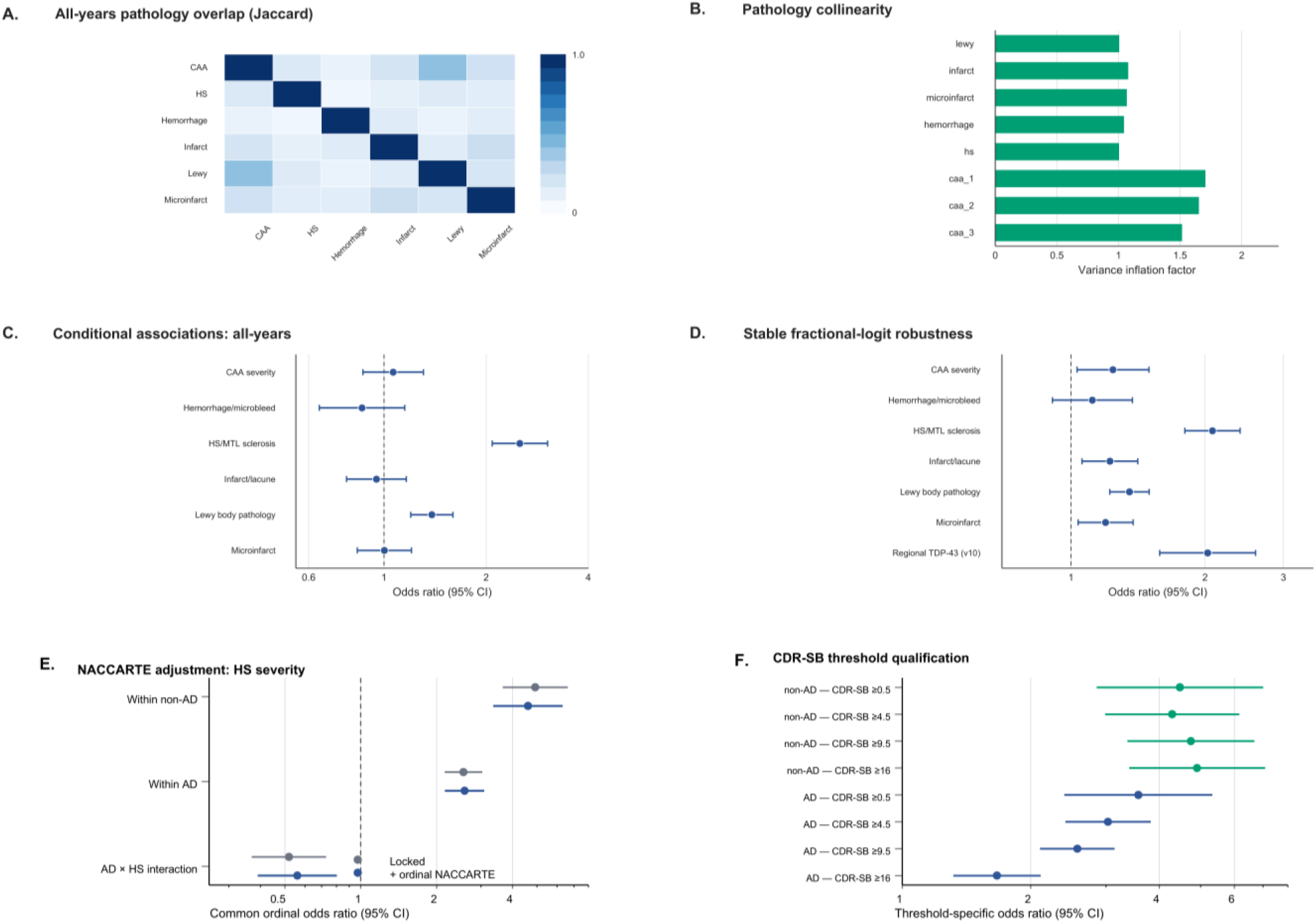
Pathology overlap and sensitivity analyses. **A)** All-years pairwise pathology overlap, summarized by Jaccard indices. **B)** Variance inflation factors for the simultaneously modeled pathology terms. **C)** Conditional associations from the all-years simultaneous model. **D)** Fractional-logit sensitivity analysis. **E)** Primary and NACCARTE-adjusted common ordinal HS estimates: non-AD, 4.93 versus 4.61; AD, 2.56 versus 2.58; and AD×HS, 0.519 versus 0.560. **F)** Threshold-specific CDR-SB associations after NACCARTE adjustment.

**Supplementary figure S3.**
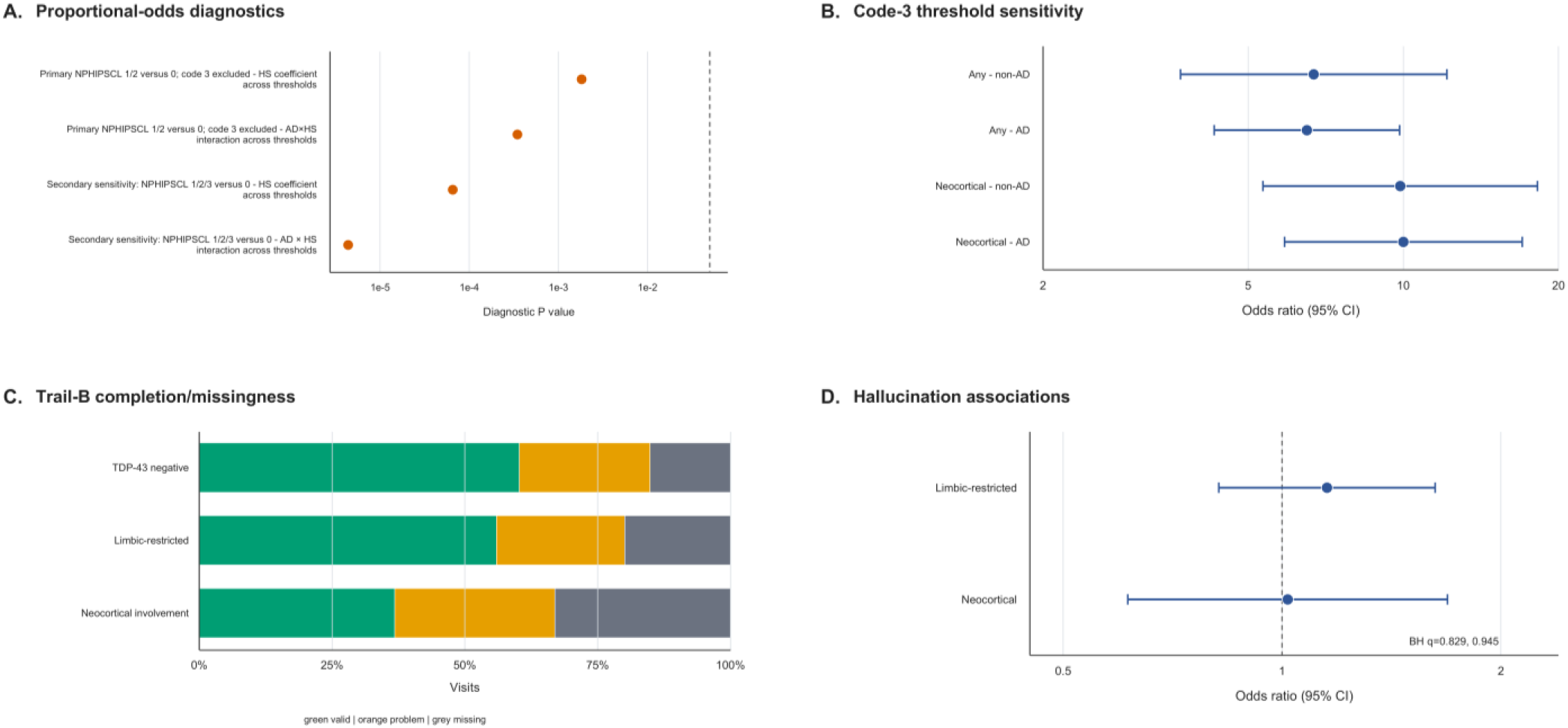
Regional TDP-43 patterns and sensitivity analyses. **A)** Proportional-odds diagnostic P values for the HS coefficient and AD×HS interaction under the primary and code-3-inclusive HS definitions; values below 0.05 indicate evidence against a common OR across thresholds. **B)** Threshold-specific HS associations when laterality-unresolved NPHIPSCL code 3 was included as HS positive. This sensitivity included 1,572 participants in the descriptive cohort and 1,529 in the covariate-complete models. **C)** Trail Making Test Part B completion, problem-coded, and missing visits by TDP-43 topography. These differences qualify the positive neocortical Trail-B estimate because test completion and missingness varied with topography and clinical severity. **D)** Adjusted hallucination associations for limbic-restricted and neocortical involvement relative to TDP-43 negativity, with BH correction across the two contrasts.

**Supplementary figure S4.**
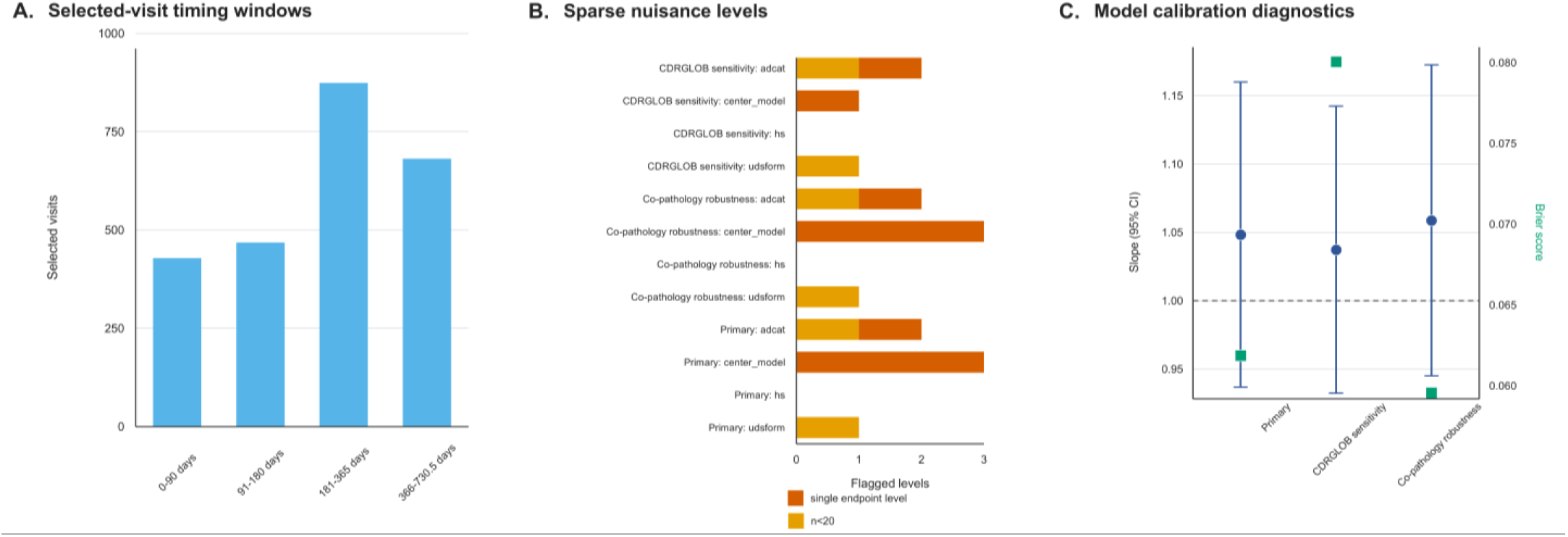
Near-death endpoint construction and model diagnostics. **A)** Distribution of selected visits across the prespecified intervals before death. **B)** Sparse levels of center, AD pathology category, HS, and UDS/form version in the primary, CDRGLOB sensitivity, and co-pathology models. **C)** Calibration slopes with 95% CIs and Brier scores for the three Firth logistic models.

**Supplementary figure S5.**
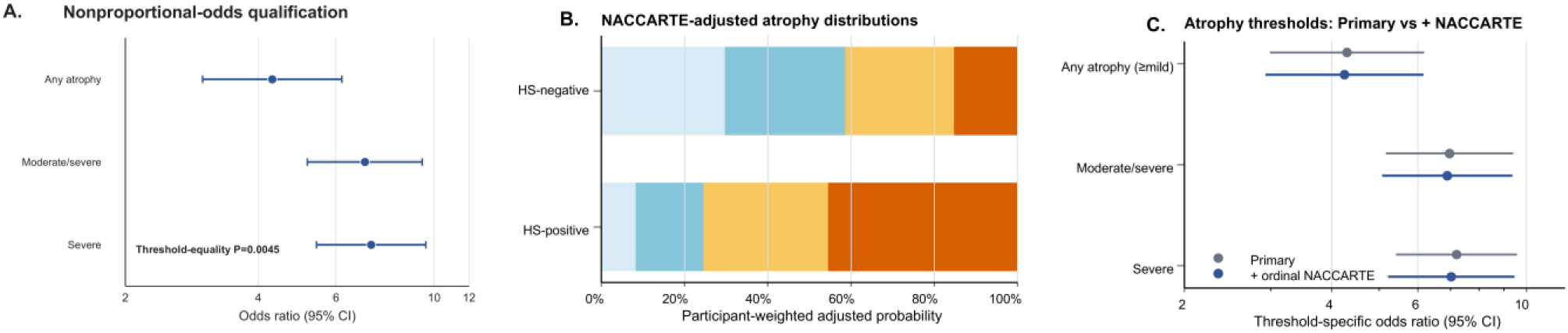
NPGRHA coverage and model diagnostics. **A)** Threshold-specific HS associations and the proportional-odds test in the primary model. **B)** Participant-weighted adjusted NPGRHA distributions after NACCARTE adjustment. **C)** Primary and NACCARTE-adjusted threshold-specific HS associations: any atrophy, OR 4.30 versus 4.25; moderate/severe atrophy, OR 6.97 versus 6.89; and severe atrophy, OR 7.20 versus 7.02.

## References

1. Hyman BT, Phelps CH, Beach TG, et al. National Institute on Aging–Alzheimer’s Association guidelines for the neuropathologic assessment of Alzheimer’s disease. Alzheimers Dement. 2012;8:1–13. doi:10.1016/j.jalz.2011.10.007.

2. Montine TJ, Phelps CH, Beach TG, et al. National Institute on Aging–Alzheimer’s Association guidelines for the neuropathologic assessment of Alzheimer’s disease: a practical approach. Acta Neuropathol. 2012;123:1–11. doi:10.1007/s00401-011-0910-3.

3. Schneider JA, Arvanitakis Z, Bang W, Bennett DA. Mixed brain pathologies account for most dementia cases in community-dwelling older persons. Neurology. 2007;69:2197–2204. doi:10.1212/01.wnl.0000271090.28148.24.

4. Boyle PA, Yu L, Leurgans SE, Wilson RS, Brookmeyer R, Schneider JA, Bennett DA. Attributable risk of Alzheimer’s dementia attributed to age-related neuropathologies. Ann Neurol. 2019;85:114–124. doi:10.1002/ana.25380.

5. Beekly DL, Ramos EM, Lee WW, et al. The National Alzheimer’s Coordinating Center (NACC) database: the Uniform Data Set. Alzheimer Dis Assoc Disord. 2007;21:249–258. doi:10.1097/WAD.0b013e318142774e.

6. Besser LM, Kukull WA, Teylan MA, et al. The revised National Alzheimer’s Coordinating Center’s Neuropathology Form—available data and new analyses. J Neuropathol Exp Neurol. 2018;77:717–726. doi:10.1093/jnen/nly049.

7. Brenowitz WD, Hubbard RA, Keene CD, et al. Mixed neuropathologies and estimated rates of clinical progression in a large autopsy sample. Alzheimers Dement. 2017;13:654–662. doi:10.1016/j.jalz.2016.09.015.

8. Amador-Ortiz C, Lin WL, Ahmed Z, et al. TDP-43 immunoreactivity in hippocampal sclerosis and Alzheimer’s disease. Ann Neurol. 2007;61:435–445. doi:10.1002/ana.21154.

9. Nelson PT, Dickson DW, Trojanowski JQ, et al. Limbic-predominant age-related TDP-43 encephalopathy (LATE): consensus working group report. Brain. 2019;142:1503–1527. doi:10.1093/brain/awz099.

10. Nag S, Yu L, Capuano AW, et al. Hippocampal sclerosis and TDP-43 pathology in aging and Alzheimer disease. Ann Neurol. 2015;77:942–952. doi:10.1002/ana.24388.

11. Gauthreaux KM, Teylan MA, Katsumata Y, et al. Limbic-predominant age-related TDP-43 encephalopathy: medical and pathologic factors associated with comorbid hippocampal sclerosis. Neurology. 2022;98:e1422–e1433. doi:10.1212/WNL.0000000000200001.

12. Woodworth DC, Nguyen KM, Sordo L, et al. Evaluating the updated LATE-NC staging criteria using data from NACC. Alzheimers Dement. 2024;20:8359–8373. doi:10.1002/alz.14262.

13. Woodworth DC, Lou JJ, Yong WH, et al. Common neuropathologic change drivers of hippocampal sclerosis of ageing. Brain. 2025;148:2400–2411. doi:10.1093/brain/awaf158.

14. Kapasi A, Yu L, Boyle PA, Barnes LL, Bennett DA, Schneider JA. Limbic-predominant age-related TDP-43 encephalopathy, ADNC pathology, and cognitive decline in aging. Neurology. 2020;95:e1951–e1962. doi:10.1212/WNL.0000000000010454.

15. Smirnov DS, Galasko D, Hansen LA, Edland SD, Brewer JB, Salmon DP. Trajectories of cognitive decline differ in hippocampal sclerosis and Alzheimer’s disease. Neurobiol Aging. 2019;75:169–177. doi:10.1016/j.neurobiolaging.2018.11.015.

16. Aiello Bowles EJ, Crane PK, Walker RL, et al. Cognitive resilience to Alzheimer’s disease pathology in the human brain. J Alzheimers Dis. 2019;68:1071–1083. doi:10.3233/JAD-180942.

17. Montine TJ, Cholerton BA, Corrada MM, et al. Concepts for brain aging: resistance, resilience, reserve, and compensation. Alzheimers Res Ther. 2019;11:22. doi:10.1186/s13195-019-0479-y.

18. Montine TJ, Corrada MM, Kawas C, et al. Association of cognition and dementia with neuropathologic changes of Alzheimer disease and other conditions in the oldest old. Neurology. 2022;99:e1067–e1078. doi:10.1212/WNL.0000000000200832.

19. Latimer CS, Burke BT, Liachko NF, et al. Resistance and resilience to Alzheimer’s disease pathology are associated with reduced cortical pTau and absence of limbic-predominant age-related TDP-43 encephalopathy in a community-based cohort. Acta Neuropathol Commun. 2019;7:91. doi:10.1186/s40478-019-0733-3.

20. Nelson PT, Lee EB, Cykowski MD, et al. LATE-NC staging in routine neuropathologic diagnosis: an update. Acta Neuropathol. 2023;145:159–173. doi:10.1007/s00401-022-02524-2.

21. Woodworth DC, Nguyen KM, Sordo L, et al. Comprehensive assessment of TDP-43 neuropathology data in the National Alzheimer’s Coordinating Center database. Acta Neuropathol. 2024;147:103. doi:10.1007/s00401-024-02728-8.

